# The complete mitochondrial genome of *Tunga penetrans* and insights into flea phylogeny

**DOI:** 10.1101/2025.06.09.658575

**Authors:** Brian Bartilol, Maureen W. Mburu, Joel Owaret, Cheryl Andisi, Isabella Oyier, Simon Muriu, Lynne Elson, Marta Maia, Joseph Mwangangi, George Githinji, Martin Rono

**Author notes:** **Corresponding authors:** Brian Bartilol & Martin Rono.

## Abstract

Tungiasis, one of the oldest and most neglected tropical diseases endemic to sub-Saharan Africa and the Americas, is caused by the female parasitic flea, *Tunga penetrans*. The flea burrows into the skin, leading to acute and chronic inflammation, often exacerbated by bacterial superinfection. Despite its significant public health impact, genomic studies on *T. penetrans* are scarce. Here, we present the first complete mitochondrial genome of *T. penetrans*, comprising 17,279 base pairs and encoding 13 protein-coding genes, 22 transfer RNAs and 2 ribosomal RNAs. Phylogenetic analysis of the cox2 gene revealed a divergent basal lineage from Brazil, supporting a South American origin of *T. penetrans* and highlights genetic differentiation within the Americas. Clustering of the Ecuadorian and African isolates further suggests historical connections, likely linked to transatlantic maritime trade. The tree highlights a strong South American origin and evidence of migration and diversification, facilitated by human and animal movement. Phylogenetic analysis of the complete genomes relying on the protein-coding genes of other fleas revealed that *T. penetrans* is closely related to *Dorcadia ioffi*, a semi-sessile flea of goats and sheep in China. This mitochondrial genome provides a critical resource for future studies on molecular epidemiology, evolutionary history, and control of tungiasis.

## Introduction

Tungiasis is one of the oldest and most neglected tropical diseases and is endemic to sub-Saharan Africa and the Americas. Tungiasis is caused by *Tunga penetrans*, one of the smallest known flea species, classified taxonomically under Phylum *Arthropoda*, Class *Insecta*, Order *Siphonaptera*, Family *Hectopsyllidae* and Genus *Tunga*. In its adult parasitic phase, the female flea burrows permanently into the human skin, while the male flea remain free-living. This is a unique adaptation, as reproduction occurs exclusively following embedment of the female flea (Nagy et al., 2007).

*T. penetrans* is thought to have originated from Central and South America before being introduced to Africa in 17^th^ century through maritime trade. It spread across Africa and the Indian continent through coastal entry points and harbours (Hoeppli, 1963). Today, an estimated 660 million people are at risk of infection, with children and the elderly disproportionately affected due to limited mobility and socioeconomic vulnerability (Deka, 2020). The disease manifests as localized inflammation at the site of female flea penetration, often complicated by secondary bacterial infections (Nyangacha et al., 2017).

In SSA, the prevalence of tungiasis is at 33%, with a higher prevalence in East Africa (34.2%) compared to West Africa (32.3%) (Obebe & Aluko, 2020). In Kenya, tungiasis represents a major public health concern, with an estimated two million people currently infected (MOH, 2014). The disease is widespread, with a heterogeneous distribution across the country. Prevalence estimates from 2021 indicated that approximately 1.3% of the population was affected by the disease (Elson, Kamau, et al., 2023). Although the distribution of the disease is heterogeneous, impoverished communities bear the greatest burden due to poor living conditions, inadequate sanitation and proximity to domestic animals (Heukelbach et al., 2001). Recent evidence shows upgrading earthen floor with affordable, cement-stabilised soil can significantly reduce the disease incidence (Elson, Nyawa, et al., 2023). Domestic animals such as pigs and dogs have also been shown to be reservoirs of tungiasis, and this underscores the need for a One Health Approach (Mutebi et al., 2015, 2016).

Despite progress in treatment strategies, the mechanism of many interventions remains poorly characterized. Furthermore, genomic data on *T. penetrans* are critically limited. Most studies have focused on single mitochondrial genes (*cox1, cox2*) or ribosomal spacers (ITS regions) (Luchetti et al., 2005, 2007). This gap hinders comprehensive insights into the parasite’s biology, transmission dynamics, and potential targets for control. To date, research on flea genomes (Order *Siphonaptera*) have prioritized mitochondrial genomes due to their compact size (15-20 kilobases), high copy number (>100) and structural conservation features that facilitate comparative and evolutionary studies.

This study addresses these gaps by presenting the first complete mitochondrial genome of *T. penetrans*, a foundational resource for advancing molecular epidemiology, understanding host-parasite coevolution, and developing targeted interventions in endemic regions (Cameron, 2025). Additionally, the knowledge on the mitochondrial genome can also inform the development of therapeutics disrupting the mitochondrial functions, a strategy that has been proven highly effective in other insects(Hao et al., 2022; Wu et al., 2016).

## Materials and methods

### Study site and ethical approval

Ethical approval was granted by the Pwani University Scientific and Ethics Review Committee (ISERC/MSc/022/2024). The study was conducted in Msabaha and Bamba villages in Kilifi County, located along the Kenya coast in Kenya. Kilifi county is an endemic area for tungiasis with an estimated prevalence of 25% (Wiese et al., 2017).

### Sample collection

Prior to flea collection, informed consent was sought from the study participants (n=10). Embedded fleas were aseptically extracted from plantar lesions using sterile 18-gauge needles under natural daylight. Each lesion was cleansed with 70% ethanol prior to extraction to minimize contamination. Fleas were carefully removed intact to preserve morphological features critical for species identification (Linardi & de Avelar, 2014). Post-extraction, participants received immediate wound care following the National Policy Guidelines on Prevention and Control of Jigger Infestations (MOH, 2014). The fleas were thereafter transported to the KEMRI-Wellcome Trust Laboratories in a cooler box at 4°C.

### Genomic DNA extraction and sequencing

Genomic Deoxyribonucleic acid (DNA) was extracted from individual fleas by crushing the fleas using a pestle and mortar and the DNA was extracted using the Qiagen Blood and Tissue kit (cat. 69506). DNA quality was assessed via a Qubit fluorometer and PCR amplification of the ribosomal intergenic spacer 2 (ITS2) region using a Q5® High-Fidelity 2X Master Mix (NEB M0492S) under published cycling conditions (Luchetti et al., 2005). PCR amplicons were resolved on a 1.5% agarose gel stained using RedSafe™ Nucleic Acid Staining Solution (iNtRON Biotechnology, Korea) and visualized using the ChemiDoc Imaging System (Bio-Rad, USA). Thereafter, the low molecular DNA was depleted from the high-molecular genomic DNA. The high molecular weight DNA was used to prepare the Nanopore Rapid sequencing genomic DNA libraries and subsequently sequenced using the Oxford Nanopore MinION platform. FASTQ reads were base called using the Guppy BaseCaller v4.0.11 from the raw FAST5 reads and stored in the FASTQ format.

### RNA extraction and sequencing

Midguts and ovaries were dissected from collected fleas for organ specific expression analysis. Total RNA was extracted from dissected tissues using the Trizol method (Rio et al., 2010). First-strand complementary DNA (cDNA) synthesis was performed using Superscript II reverse transcriptase (Invitrogen, cat. 18064014) and oligo (dT) primers (Invitrogen, cat. AM1710), followed by second strand synthesis using the Klenow fragment (New England Biolabs, Ipswich, US, cat. M0210S). The double-stranded DNA was normalized to 10 ng/µL as part of quality control protocols prior to library preparation. Sequencing libraries were constructed using the Nextera library preparation kit (Illumina, San Diego, US, cat. FC-121-1031), which employs a tagmentation step to fragment DNA and ligate adapters simultaneously. PCR amplification was then conducted to enrich libraries and incorporate dual-indexing barcodes. To ensure library integrity, fragment size distribution and quality were assessed using a Bioanalyzer 2100 (Agilent Technologies, Santa Clara, US), confirming a target range of 300–700 bp suitable for sequencing. Libraries were sequenced on an Illumina MiSeq platform with 150 bp paired-end chemistry.

### Assembly, polishing and annotation

Long-read sequencing data were initially mapped to the mitochondrial genome of *Dorcadia ioffi*, a close relative of *T. penetrans* and aligned reads were extracted. This step enriches mitochondrial genome sequences and filters out nuclear genomic DNA. *De Novo Assembly* was performed using Canu v2.2 with parameters optimized for flea mitochondrial genome based on other flea mitochondrial genome sizes and nanopore data (genomeSize=16k, -nanopore-raw) (Koren et al., 2017). The resulting assembly was polished using Illumina short reads with Pilon (Walker et al., 2014), employing the --fix all option to resolve SNPs and indels. (Walker et al., 2014). The long and short reads were mapped back to the genome using HISAT2 v2.2.1 and Minimap2 v2.24 respectively(Kim et al., 2019; Li, 2018). The resulting alignments were converted into sorted and indexed BAM format using SAMtools v1.15.1 and per-base coverage was calculated using Bedtools v2.30.0 (Danecek et al., 2021; Quinlan & Hall, 2010). The output was then exported in a CSV format and visualized in R software(R Core Team, 2024).

Mitogenome annotation was conducted using Mitos2 (Bernt et al., 2013) with the invertebrate genetic code and the refseq89o reference database (Bernt et al., 2013). Open reading frames (ORFs) for protein-coding genes were manually curated in Geneious Prime® 2025.1.1 by comparative alignment with mitochondrial genomes of *Dorcadia ioffi* and *Pulex irritans*. The final mitochondrial genome was deposited in GenBank under accession number PV426769 and visualized as a circular map using the *circularMT* tool (Goodman & Carr, 2024)

### Phylogenetic analysis

Protein-coding mitochondrial sequences of fleas (Order Siphonaptera) were retrieved from GenBank, with *Casmara patrona* (Order Lepidoptera) included as an outgroup (Supplementary Table 1) (Jiang et al., 2021a). Sequences (n=9) were aligned using Muscle v5.3 (Edgar, 2004) and phylogenetic reconstruction carried out using IQ-TREE v 2.4.0 (Nguyen et al., 2014). The analysis employed the *-m* TEST and -*b* 1000 parameters to automatically select the best-fit substitution model and perform 1000 ultrafast bootstraps replicates, respectively (Nguyen et al., 2014). The best model in this case was mtART+F+I+G4 according to the Bayesian Information Criterion (BIC). The resulting tree was visualized and annotated using the ggtree package (Yu et al., 2017) in R software (R Core Team, 2024). For intraspecific analysis of *T. penetrans*, 16 globally distributed mitochondrial sequences of the cox2 gene (Supplementary Table 2) were downloaded from GenBank (Luchetti et al., 2007). Alignment and phylogenetic inference followed the same workflow (Muscle alignment, IQ-TREE parameters, and *ggtree* visualization) (R Core Team, 2024; Yu et al., 2017). The best model selected in this case was HKY+F according to BIC.

**Table 1.**
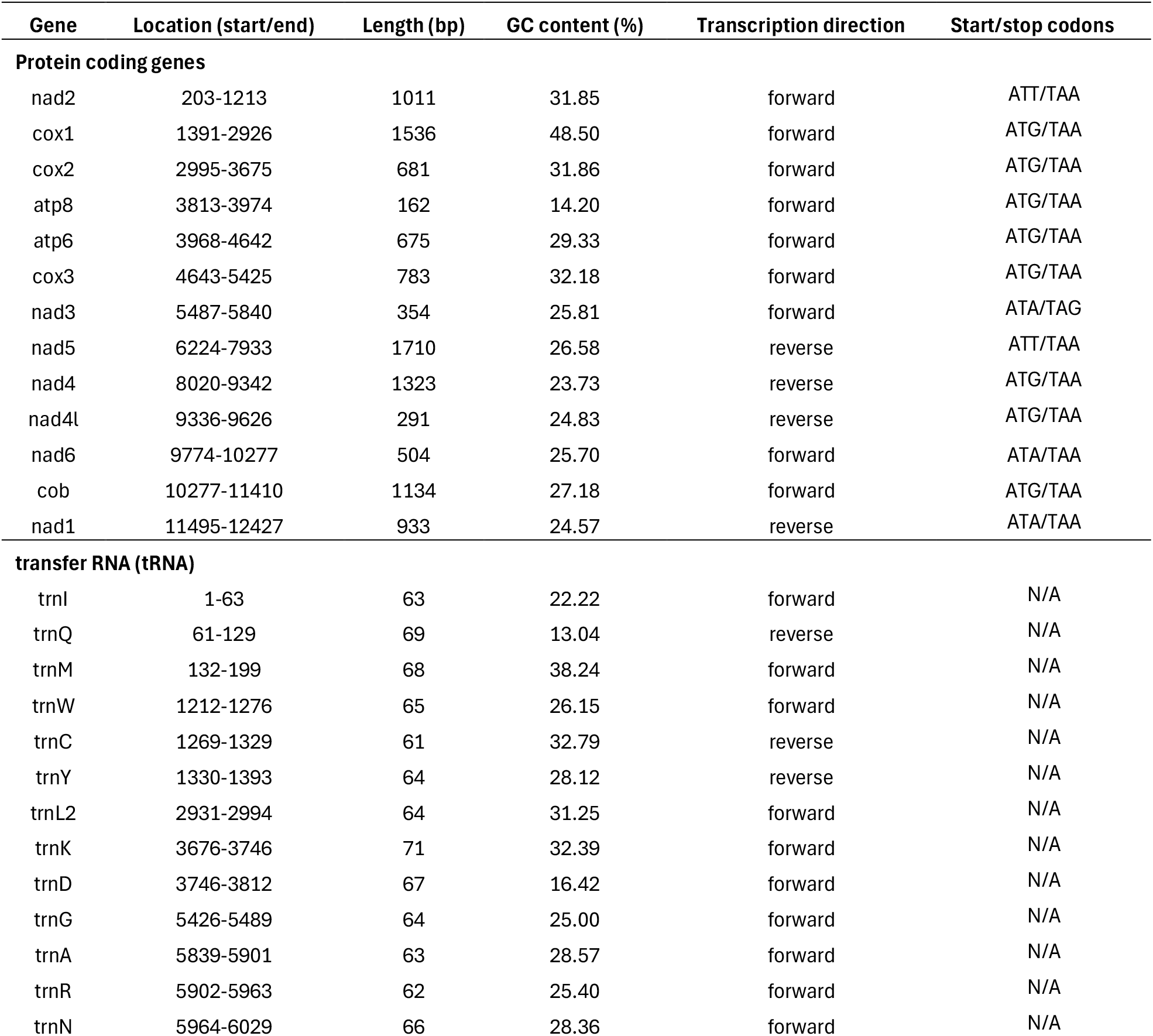
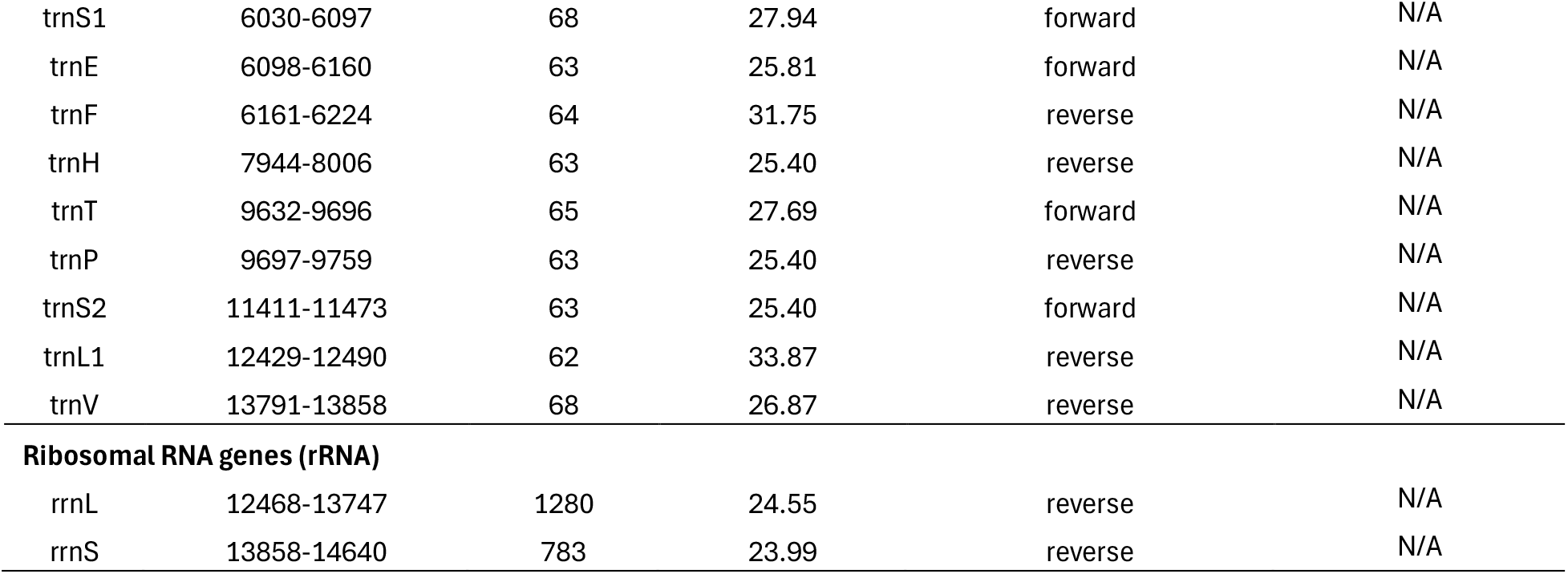
The mitochondrial genes of T. penetrans and their location, lengths, GC contents, direction of transcription and the start/stop codons.

## Results and Discussion

### Characteristics of the *Tunga penetrans* mitochondrial genome

The mitochondrial genome of *T. penetrans* spans 17,279 base pairs long and encodes 37 genes, including 13 protein-coding genes (PCGs), 22 transfer RNA (tRNA) genes, and two ribosomal RNA (rRNA) genes. The PCGs comprise subunits of key respiratory complexes: *atp6* and *atp8* (ATP synthase), *cox1*-*cox3* (cytochrome c oxidase), *cytb* (cytochrome b), and *nad1*–*nad6* and *nad4l* (NADH dehydrogenase). The 22 tRNAs, covering all canonical amino acids, are designated as *trnI, trnQ, trnM, trnW, trnC, trnY, trnL2, trnK, trnD, trnG, trnA, trnR, trnN, trnS1, trnE, trnF, trnH, trnT, trnP, trnS2, trnL1*, and *trnV*. The genome also included the small (*rrnS)* and large (*rrnL*) ribosomal RNA subunits, along with a replication origin (Figure 1a). The nucleotide composition is heavily AT-biased (A: 40.2%, T: 41.8%), with lower GC content (C: 10.59%, G:7.41%) resulting in a total AT content of 82%, consistent with other flea mitogenomes (Table 1). Among PCGs, cox3 (32.18%), cox2 (31.86%) and nad2 (31.85%) exhibited the highest GC content, suggesting potential selective pressure or structural constraints. RNA-seq read mapping revealed elevated coverage in the large ribosomal RNA subunit (*rrnL*) and cytochrome c oxidase genes (cox1-cox3), likely reflecting their high transcription activity in the mitochondrial energy production (Figure 1b).

**Figure 1.**
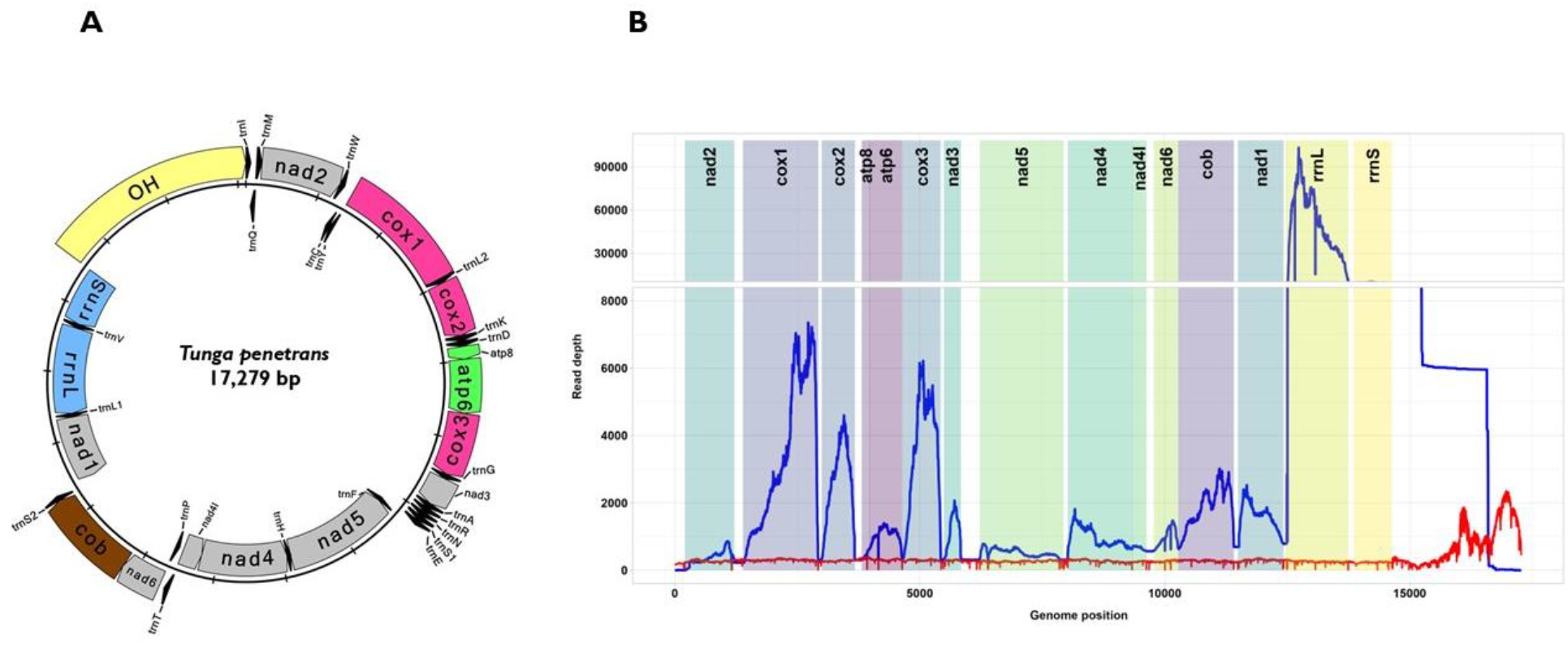
Mitochondrial genome organization and transcriptional activity in *T. penetrans*. A) A circular plot depicting the genomic organization of the complete mitochondrial genome of human parasite, *Tunga penetrans*. The mitochondrial genome encodes 13 protein coding genes, 22 transfer RNAs and 2 ribosomal RNA genes, along with a replication origin. Genes are color-coded by functional category and transcribed in clockwise (outer circle) or counterclockwise (inner circle) directions. The genome’s organization and gene order are conserved relative to other Siphonaptera species. B) RNA-Seq read coverage across the mitochondrial genome of *T. penetrans*. The coverage plot highlights regions of high transcriptional activity, particularly in the large ribosomal RNA subunit (*rrnL*) and cytochrome c oxidase genes (*cox1-cox3*). RNA-Seq reads were mapped to the mitochondrial genome to validate and refine the assembly, ensuring accuracy in gene annotation. The red line indicates number of sequences from the nanopore reads that mapped back to the genome.

### Relationship of Tunga penetrans with other fleas of Siphonaptera order

The maximum-likelihood phylogeny resolves nine flea species (Order Siphonaptera) into a monophyletic group, with the moth *Casmara patrona* (Lepidoptera) positioned as the outgroup (Figure 2). Within Siphonaptera, the members of the family Pulicidae form a well-supported and distinct clade. Earlier diverging lineages include members of the families Ceratophyllidae (*Jellisonia amadoi, Ceratophyllus wui*), Hystrichopsyllidae (*Hystrichopsylla weida*), Vermipsyllidae (*Dorcadia ioffii*) and Hectopsyllidae (*T. penetrans*), which occupy more basal positions relative to Pulicidae. Notably, *T. penetrans* (Hectopsyllidae) cluster more closely with Dorcadia *ioffi* (Vermipsyllidae), supporting the sister-group relationship between these two fleas.

**Figure 2.**
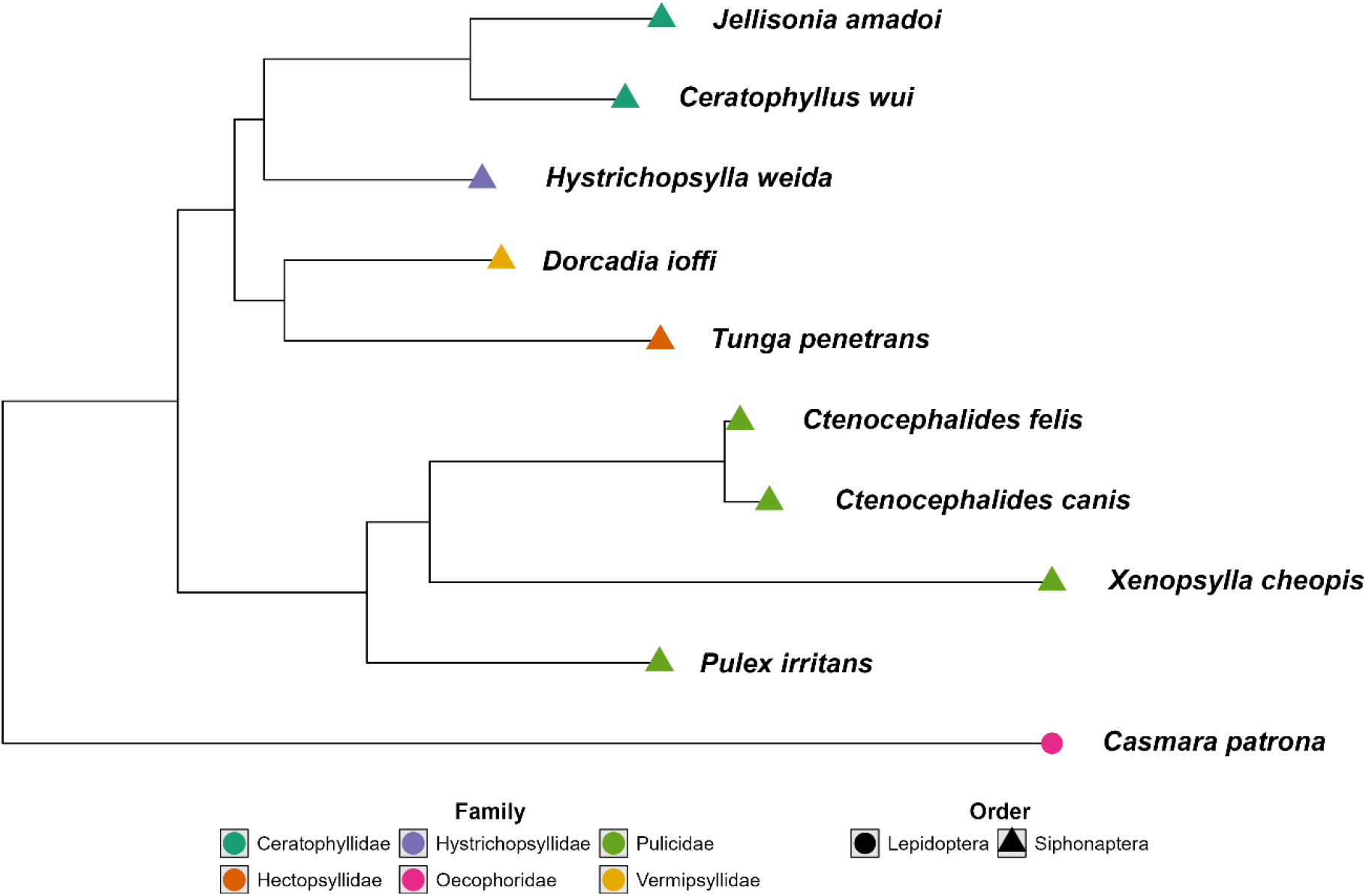
Maximum-likelihood phylogenetic tree of complete mitochondrial genomes of the members of the order *Siphonaptera*, rooted with *Casmara patrona* (Lepidoptera) as an outgroup. The mitochondrial genome of *T. penetrans* (Hectopsyllidae) clusters more closely related with *Dorcadia ioffi* (Vermipsyllidae), indicating a sister-group relationship.

### Relationship with other *Tunga penetrans* sequenced globally

The phylogenetic tree on cox2 gene of isolates from Brazil, Burundi, Democratic Republic of the Congo, Ecuador, Kenya, and Madagascar revealed a divergent basal lineage from Brazil (Figure 3). The other isolates clustered into a more recent and diverse clade with included isolates from Burundi, Democratic Republic of the Congo, Ecuador, Kenya, and Madagascar. Within this subclade, a distinct subclade was observed consisting primarily of Ecuadorian isolates. The rest of the clade contained Ecuadorian isolates interspersed with isolates from African countries.

**Figure 3.**
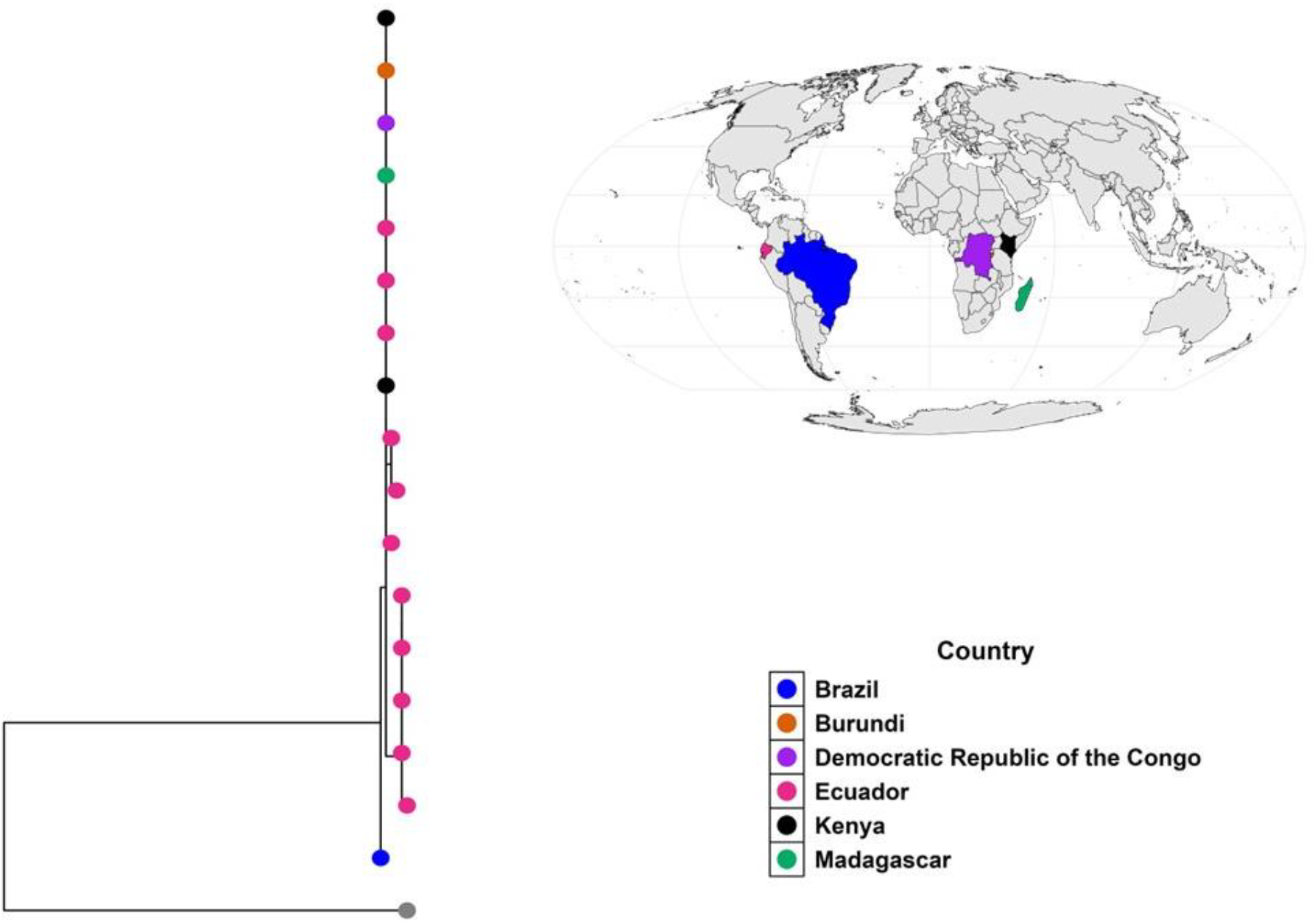
The phylogenetic relationship of *T. penetrans* collected globally based on the *cox2* gene. The tree resolves two distinct clades: one exclusively comprising Ecuadorian haplotypes and another encompassing isolates from Brazil, Burundi, the Democratic Republic of the Congo (DRC), Kenya, and Madagascar.

## Discussion

Here, we present the first complete mitochondrial genome assembly of *T. penetrans*, a flea responsible for significant morbidity in both humans and domestic animals. The genome spans 17,279 bp and exhibits a canonical metazoan mitochondrial structure, encoding 13 PCGs, 22 tRNAs, and 2 rRNA genes, and replication origin. The genome displays pronounced AT bias (82% AT content) consistent with other characterized flea mitogenomes within the order Siphonaptera (Cameron, 2015; Tan et al., 2023; Verhoeve et al., 2020; Xiang et al., 2017). Among PCGs, *cox3, cox2*, and *nad2*, exhibited the highest GC content, likely reflecting structural or functional constraints in these essential oxidative phosphorylation components. RNA-seq analysis revealed elevated transcriptional activity in the large ribosomal RNA subunit and cytochrome c oxidase genes, underscoring their central roles in mitochondrial energy production and ribosome biogenesis.

Phylogenetic analysis using complete mitochondrial genomes of Siphonaptera species resolved *T. penetrans* (Hectopsyllidae family) as closely related to *Dorcadia ioffi*, super-family of Vermipsyllidae *(Cameron, 2015; Tan et al., 2023; Verhoeve et al., 2020; Xiang et al., 2017; Zhang et al., 2022). Dorcadia ioffi* is a semi-sessile flea of goats and sheep in China, Mongolia and Russia (Rothschild, 1992; Xiang et al., 2017). The close relationship is also corroborated by several shared morphological and physiological changes. Both *T. penetrans* and *D. ioffi* exhibit sessile tendencies with varying degrees on their host. They also undergo dramatic neosomy, which is a post mating hypertrophy that transforms the female’s abdomen without molting to accommodate the large number of eggs. Additionally, the males of both species mate with the attached females in situ on the host(Dowling, 2015; Rothschild, 1992). These finding advances our understanding of flea evolutionary relationships and corroborate the monophyly of the Siphonaptera, with *Casmara patrona* (Lepidoptera) as an outgroup(Jiang et al., 2021b). Within this order, the family Pulicidae formed a distinct, derived clade, while families Ceratophyllidae, Hystrichopsyllidae, and Vermipsyllidae diverged earlier, aligning with prior phylogenetic hypothesis (Zhang et al., 2022). Although the complete mitochondrial genomes offer this resolution within Siphonaptera, whole nuclear genomes could provide a finer resolution.

The phylogenetic analysis of the cox2 gene reveals an ancestral from Brazil supporting the hypothesis for a South American origin for *T. penetrans* and indicating early evolutionary separation from other isolates. Within the main clade, most Ecuadorian isolates clustered together suggesting geographical clustering, while others interspersed with African isolates from Kenya, Burundi, the Democratic Republic of the Congo, and Madagascar. This points to multiple introduction events or historical gene flow between South America and Africa, likely driven by transatlantic maritime trade and human or animal movement. One such evidence supporting this theory includes various reports documenting imported cases of tungiasis from endemic to non-endemic regions worldwide including China, Greece, Japan, Portugal, Taiwan, Unites States and the United Kingdom (Chen et al., 2011; Douvali et al., 2025; Hager et al., 2008; Hakeem et al., 2010; Santos et al., 2017; Yotsu et al., 2011). Furthermore, the distinct Ecuadorian subclade along with dispersed isolates suggests the presence of structured populations in Ecuador, and this was attributed to geographical isolation of some of the collection sites(Luchetti et al., 2005). The lack of distinct clustering among African isolates could be due to shared ancestry or regional genetic mixing within the continent. While this data reveals Brazilian ancestral lineage and potential historical transmission pathways, the direction of the gene flow cannot be conclusively be inferred from mitochondrial genome data alone.

## Conclusion

This study presents the first complete mitochondrial genome of *T. penetrans*, a critical resource for understanding its genomic architecture, evolutionary history, and its global phylogeography. The genome AT-rich composition (82%) and conserved gene order align with features observed in other Siphonaptera species, reinforcing shared evolutionary constraints within the order. Phylogenetic analysis of the complete mitochondrial genomes showed that *T. penetrans* is closely related to D. ioffii, a flea of goats and sheep. This can further be corroborated by the shared morphological and physiological characteristics such as sessility (though its intensity varies), neosomy in females and in situ mating by males. Phylogenetic analysis of the cox2 gene supports a South American origin of T. penetrans, with a Brazilian ancestral origin and early divergence from other global isolates. Ecuadorian samples showed evidence of geographical clustering, while African isolates appeared more genetically mixed, pointing to recent intercontinental dispersal. These patterns suggest multiple introduction events and historical genetic flow between South America and Africa, likely facilitated by transatlantic trade and human and animal movement. Although these findings reveal potential transmission pathways, the direction of gene flow cannot be determined from mitochondrial genome data alone.

This mitochondrial genome provides a foundational tool for future studies on *T. penetrans* adaptation mechanisms, population genetics, and targeted control strategies in endemic regions. By elucidating evolutionary trajectory and genetic diversity, this work advances efforts to mitigate tungiasis-related morbidity in vulnerable populations. Notably, in agricultural pest management, mitochondrial genomes of insects have successfully been targeted using chemical agents and RNA interference technologies, highlighting the potential for similar approaches in *T. penetrans* control.

## Acknowledgement

I would like to acknowledge the study participants for allowing us to collect the jigger samples and Ken Muriithi who assisted with sample collection.

## Ethical statement

This study received ethical approval from the Pwani University Ethical Review Committee. Prior to collecting jiggers (*T. penetrans*) from participants, informed consent was obtained from parents/guardians, and assent was secured from children aged 12 years and older. Following sample collection, all participants and affected community members received treatment for tungiasis in accordance with the National Policy Guidelines on Prevention and Control of Jigger Infestations.

## Data Accessibility

The complete mitochondrial genome of *Tunga penetrans* has been deposited in the GenBank database under the accession number PV426769.1. All supporting data and R scripts are available at https://github.com/bartilol/Tunga-penetrans-mitochondial-genome-data-and-scripts.git. Additional data may also be found in the supplementary files.

