## Supplementary material for "The complete mitochondrial genome of *Tunga penetrans* and insights into flea phylogeny": Supplemenentary Table 1

Supplementary Table 1. The accession number of the complete mitochondrial genomes of the Order Siphonaptera

| **Accession no.** | **Definition** |
|---|---|
| MW310242.1 | *Xenopsylla cheopis* mitochondrion, complete genome |
| NC_022710.1 | *Jellisonia amadoi* mitochondrion, complete genome |
| NC_036066.1 | *Dorcadia ioffi* mitochondrion, complete genome |
| NC_040301.1 | *Ceratophyllus wui* mitochondrion, complete genome |
| NC_042380.1 | *Hystrichopsylla weida* qinlingensis mitochondrion, complete genome |
| NC_049858.1 | *Ctenocephalides felis* isolate EL2017-DRISC mitochondrion, complete genome |
| NC_063709.1 | *Pulex irritans* mitochondrion, complete genome |
| NC_063710.1 | *Ctenocephalides canis* mitochondrion, complete genome |
| PV426769.1 | *Tunga penetrans* mitochondrion, complete genome |
