## Supplementary material for "The complete mitochondrial genome of *Tunga penetrans* and insights into flea phylogeny": Supplemenentary Table 2

Supplementary Table 2. Details on the cox2 sequences used for the intraspecific phylogenetic analysis using the cox2 gene sequences

| Accession no | Country | Location |
|---|---|---|
| AY425821.1 | Ecuador | S.ta Isabel |
| DQ844701.1 | Ecuador | Ambuqui |
| AF551754.1 | Ecuador | S.ta Isabel (Site B) |
| DQ844698.1 | Ecuador | Squisil |
| AF551752.1 | Ecuador | Pelileo |
| DQ844706.1 | Ecuador | S. Francisco de las Pampas |
| DQ844705.1 | Ecuador | S. Francisco de las Pampas |
| DQ844704.1 | Ecuador | S. Francisco de las Pampas |
| DQ844695.1 | Kenya | Wamba |
| DQ844703.1 | Ecuador | S. Francisco de las Pampas |
| DQ844699.1 | Ecuador | Olmedo |
| AF551753.1 | Madagascar | |
| DQ844697.1 | Burundi | Mivo |
| DQ844696.1 | Democratic Republic of the Congo | Kangu |
| PV426769.1 | Kenya | Kilifi |
| DQ844702.1 | Ecuador | Parambas |
| DQ844700.1 | Brazil | Fortaleza |
